# Functional testing of a human *PBX3* variant in zebrafish reveals a potential modifier role in congenital heart defects

**DOI:** 10.1101/337832

**Authors:** Gist H. Farr, Kimia Imani, Darren Pouv, Lisa Maves

## Abstract

Whole-genome and whole-exome sequencing efforts are increasingly identifying candidate genetic variants associated with human disease. However, predicting and testing the pathogenicity of a genetic variant remains challenging. Genome editing allows for the rigorous functional testing of human genetic variants in animal models. Congenital heart defects (CHDs) are a prominent example of a human disorder with complex genetics. An inherited sequence variant in the human *PBX3* gene (*PBX3* p.A136V) has previously been shown to be enriched in a CHD patient cohort, indicating that the *PBX3* p.A136V variant could be a modifier allele for CHDs. *PBX* genes encode TALE (Three Amino acid Loop Extension)-class homeodomain-containing DNA-binding proteins with diverse roles in development and disease and are required for heart development in mouse and zebrafish. Here we use CRISPR-Cas9 genome editing to directly test whether this *PBX* gene variant acts as a genetic modifier in zebrafish heart development. We used a single-stranded oligodeoxynucleotide to precisely introduce the human *PBX3* p.A136V variant in the homologous zebrafish *pbx4* gene (*pbx4* p.A131V). We find that zebrafish that are homozygous for *pbx4* p.A131V are viable as adults. However, we show that the *pbx4* p.A131V variant enhances the embryonic cardiac morphogenesis phenotype caused by loss of the known cardiac specification factor, Hand2. Our study is the first example of using precision genome editing in zebrafish to demonstrate a function for a human disease-associated single nucleotide variant of unknown significance. Our work underscores the importance of testing the roles of inherited variants, not just *de novo* variants, as genetic modifiers of CHDs. Our study provides a novel approach toward advancing our understanding of the complex genetics of CHDs.

**Summary statement:** Our study provides a novel example of using genome editing in zebrafish to demonstrate how a human DNA sequence variant of unknown significance may contribute to the complex genetics of congenital heart defects.

## Introduction

Whole-genome and whole-exome sequencing efforts are increasingly identifying genetic variation in the general human population as well as candidate genetic variants associated with disease (Lek et al., 2016; Wright et al., 2015). There are several approaches for predicting and testing the pathogenicity of a genetic variant (Cox et al., 2015; Richards et al., 2015; Starita et al., 2017). However, demonstrating a function of a particular sequence variant remains challenging. Genome editing now allows for the precise engineering of human genetic variants in animal models for rigorous functional testing (Doudna and Charpentier, 2014; Peng et al., 2014). There are examples of the use of CRISPR-Cas9 engineering in mouse and *C. elegans* animal models to demonstrate functional effects of human disease-associated sequence variants (Arno et al., 2016; DiStasio et al., 2017; Lin et al., 2016; Prior et al., 2017). However, it is not yet clear how effectively human variants of unknown significance can be functionally tested through genome editing in animal models.

Congenital heart defects (CHDs) are a prominent example of a human disorder with complex genetics (reviewed in Fahed et al., 2013; Gelb and Chung, 2014; Zaidi and Brueckner, 2017). CHDs occur in about 1% of live births and are the leading cause of infant death due to birth defects. Intensive studies have uncovered prominent roles for transcription and chromatin factors in heart development and CHDs (Chang and Bruneau, 2011; Fahed et al., 2013; Gelb and Chung, 2014). Cardiac transcription factors of the GATA, HAND, MEF2, NKX, SRF, and TBX families are required for heart development in mice and zebrafish animal models, and mutations in genes encoding these factors can cause human CHDs (Evans et al., 2010; McCulley and Black, 2012; Olson, 2006). Large-scale whole-exome sequencing studies find that CHD cases show an excess of *de novo* mutations for many transcription and chromatin factors (Homsy et al., 2015; Zaidi et al., 2013). In spite of these efforts, these *de novo* mutations likely account for only about 10% of CHDs (Gelb and Chung, 2014; Homsy et al., 2015; Zaidi et al., 2013). While additional studies are identifying potential contributions of inherited mutations in CHDs (for example, Jin et al., 2017), our understanding of the genetics of CHDs is still incomplete.

The genetics of CHDs is complex, in part, because the same candidate gene, and even the same sequence variant, can be associated with a spectrum of heart malformations and can even be present in control cases (reviewed in Fahed et al., 2013; Gelb and Chung, 2014; Zaidi and Brueckner, 2017). This is exemplified in studies of *NKX2.5* mutations in CHD patients and families (Elliott et al., 2003; McElhinney et al., 2003; Stallmeyer et al., 2010). Thus, genetic risk factors, or modifier genes, likely influence the phenotype of CHDs, but modifier genes are difficult to identify and characterize (Fahed et al., 2013; Gelb and Chung, 2014; Zaidi and Brueckner, 2017). In order to understand the etiology of CHDs and the roles of genetic risk factors in influencing the phenotypic spectrum of CHDs, it is imperative that we increase our understanding of how genetic variants and modifier alleles regulate heart development and contribute to CHDs.

Sequence variants in human *PBX* genes have been identified in patients with CHDs (Arrington et al., 2012; Slavotinek et al., 2017), indicating that these *PBX* variants could contribute to CHDs. One of these variants, an inherited missense variant in the coding region of the *PBX3* gene (p.A136V; 9:128678097 C>T), occurred at a frequency of 2.6% in a cohort of CHD patients (0.66% in controls; Arrington et al., 2012). These patients exhibited a spectrum of CHDs, particularly outflow tract malformations (Arrington et al., 2012). *Pbx* genes encode TALE (Three Amino acid Loop Extension)-class homeodomain-containing DNA-binding proteins, which have diverse roles in development and disease (Cerdá-Esteban and Spagnoli, 2014; Moens and Selleri, 2006). In mouse embryos, *Pbx1* is required for heart development, and loss of varying alleles of *Pbx1, Pbx2, and Pbx3* leads to a spectrum of cardiac outflow tract defects (Chang et al., 2008; Stankunas et al., 2008). Our previous studies have shown that zebrafish Pbx proteins are also needed for outflow tract development and are needed for early myocardial differentiation and morphogenesis (Kao et al., 2015; Maves et al., 2009). The *PBX3* p.A136V variant lies in a highly conserved poly-Alanine tract, which has been implicated in Pbx binding to histone deacetylase (HDAC) chromatin proteins (Saleh et al., 2000). While this variant is predicted to be deleterious (Arrington et al., 2012), it is not known whether it affects the function of *PBX3* or contributes to CHD. Because this variant is present in controls and has the potential to be inherited, it may represent a modifier or risk factor for CHDs. The *PBX3* p.A131V variant is present in the human population with an allele frequency of >0.6% (Arrington et al., 2012; ExAC, 2018) and so is likely excluded from studies of *de novo* or rare inherited variants associated with CHDs (Homsy et al., 2015; Jin et al., 2017; Sifrim et al., 2016; Zaidi et al., 2013).

Here we use CRISPR-Cas9 genome editing in zebrafish to test whether the *PBX3* p.A131V variant could function as a modifier allele in CHDs. In particular, we test whether this variant enhances the phenotype caused by loss of a cardiac specification factor, Hand2, in zebrafish heart development. Our study is the first example, of which we are aware, of using precision genome editing in zebrafish to demonstrate a function for a human disease-associated single nucleotide variant of unknown significance. Our work underscores the importance of testing the roles of inherited variants as genetic modifiers of CHDs.

## Results

### Zebrafish *pbx4*, but not *pbx3b*, is required for early cardiac morphogenesis

An inherited heterozygous variant in *PBX3*, c.407C>T, predicting p.Arg136>Val, was previously identified as enriched in a cohort of patients with congenital heart defects (Arrington et al., 2012). The allelic frequency of this variant was significantly less frequent in the study’s control population (P = 0.047; Arrington et al., 2012) and is also significantly less frequent in the current ExAC database population (P = 0.0047, Chi-square test; ExAC, 2016). This *PBX3* p.A136V variant is present in a highly conserved poly-Alanine tract, which lies between the two PBC domains that interact with HDAC and Meis proteins (Fig. 1A; Choe et al., 2009; Saleh et al., 2000). The human and zebrafish Pbx genes that do not show 100% conservation of the poly-Alanine tract (human *PBX4* and zebrafish *pbx2* and *pbx3a*; Fig. 1A) appear to have reduced functional roles in development, as human *PBX4* variants are observed at near expected frequencies in the general population (ExAC, 2018) and zebrafish *pbx2* null mutants are homozygous viable as adults (our unpublished data). However, loss of function variants in *PBX3* are highly underrepresented in the ExAC database (ExAC, 2018), suggesting that human *PBX3* is required for viability. The enrichment of the *PBX3* p.A136V variant in a CHD patient cohort suggests that it may contribute a modifier role in the complex genetics of congenital heart defects and led us to investigate a potential function of this allele in zebrafish.

**Figure 1.**
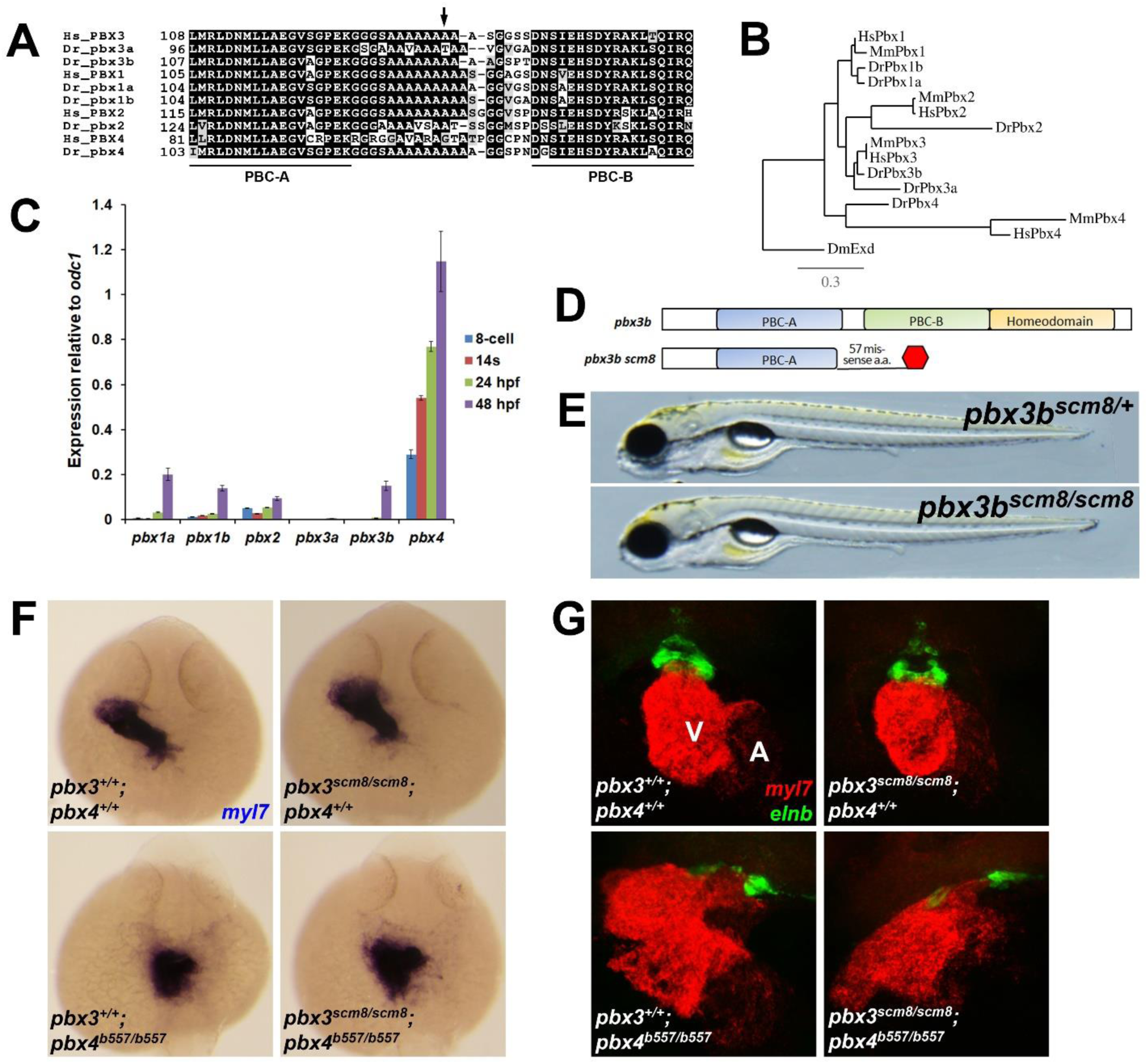
Zebrafish *pbx4*, but not *pbx3b*, is required for early cardiac morphogenesis. (A) Alignments of human (Hs) and zebrafish (Dr) Pbx proteins in the region of the poly-Alanine tract. Numbers indicate amino acid positions. Partial PBC-A and PBC-B domains are underlined. Arrow marks position of amino acid 136 in human PBX3. (B) Phylogenetic analysis of human (Hs), mouse (Mm), and zebrafish (Dr) Pbx genes. DmExd is the Drosophila Pbx ortholog Extradenticle. (C) qRT-PCR analysis of *pbx* gene expression in wild-type zebrafish embryos at four embryonic stages: 8-cell (about 1.25 hours post fertilization, hpf), 14 somites (s; about 16 hpf), 24 hpf, and 48 hpf. Levels of expression each *pbx* gene are shown relative to the expression of *odc1*. Error bars represent standard deviations for three technical replicates. (D) Schematic of zebrafish Pbx3b protein domains and inferred domains encoded by the CRISPR-Cas9-generated *pbx3b*^*scm8*^ allele. (E) Images of live, 5 days post fertilization (dpf) *pbx3b*^*scm8/+*^ and *pbx3b*^*scm8/scm8*^ larvae. *pbx3b*^*scm8/scm8*^ larvae show no obvious heart or other defects at least up to 7 dpf. (F) Myocardial marker *myl7* expression at 24 hpf appears normal in *pbx3*^*+/+*^ *;pbx4*^*+/+*^ (n = 11), in *pbx3*^*scm8/scm8*^*;pbx4*^*+/+*^ (n = 2), and in *pbx3*^*scm8/scm8*^*;pbx4*^*b557/+*^ (n = 15) embryos. *pbx3*^*+/+*^ *;pbx4*^*b557/b557*^ (n = 8) and *pbx3*^*scm8/scm8*^*;pbx4*^*b557/b557*^ (n = 7) embryos show similarly disrupted early heart tube morphogenesis, swe previously described for *pbx4*^*b557/b557*^ embryos (Kao et al., 2015). Dorsal views; anterior is up. (G) Expression of myocardial marker *myl7* (red) and outflow tract marker *elnb* (green) at 60 hpf appears normal in *pbx3*^*+/+*^ *;pbx4*^*+/+*^ (n = 8) and in *pbx3*^*scm8/scm8*^*;pbx4*^*+/+*^ (n = 9) embryos. *pbx3*^*+/+*^ *;pbx4*^*b557/b557*^ (n = 11) and *pbx3* ^*scm8/scm8*^*;pbx4*^*b557/b557*^ (n = 10) embryos show variably disrupted myocardial and outflow tract morphogenesis, similar to what we previously described for *pbx4*^*b557*^*/*^*b557*^ embryos (Kao et al., 2015). V, ventricle. A, atrium. Ventral views, anterior is up.

In zebrafish and in mice, multiple *Pbx* genes are expressed broadly and have high functional redundancy (Capellini et al., 2004; Ferretti et al., 2011; Moens and Selleri, 2004; Pöpperl et al., 2000; Ruzicka et al., 2015). Zebrafish have six *pbx* genes, and *pbx3b* is the closest ortholog of human *PBX3* (Fig. 1B). Neither *pbx3b* nor *pbx3a* are detectably expressed during early zebrafish development prior to about 48 hpf (hours post fertilization; Fig. 1C; Ruzicka et al., 2015; Waskiewicz et al., 2002). Because *pbx3b* has a conserved poly-Alanine tract (Fig. 1A) and shows upregulated expression by 48 hpf (Fig. 1C) (after heart tube formation in zebrafish but around the stage of early outflow tract development; Grimes et al., 2006), we tested the function of *pbx3b* by using CRISPR-Cas9 to generate zebrafish *pbx3b* mutants. The *pbx3b*^*scm8*^ mutation that we generated creates an early stop codon (Fig. 1D). We find that zebrafish *pbx3b*^*scm8/scm8*^ larvae are viable and appear normal (Fig. 1E) and that *pbx3b* does not show any detectable requirements during early heart development, as heart tube formation, heart chamber formation, and outflow tract development appear grossly normal in *pbx3b*^*scm8/scm8*^ embryos (Fig. 1F-G). *pbx4*, however, is the main zebrafish *pbx* gene expressed during early zebrafish development (Fig. 1C; Waskiewicz et al., 2002). We have previously shown that zebrafish *pbx4* is needed for proper heart and outflow tract development (Kao et al., 2015; Maves et al., 2009). In particular, *pbx4* mutant embryos show disrupted myocardial morphogenesis and variable defects in outflow tract development (Kao et al., 2015; Fig. 1F-G). Because Pbx genes have been shown to function redundantly (Capellini et al., 2006; Ferretti et al., 2011; Maves et al., 2007; Waskiewicz et al., 2002), we tested whether *pbx3b* and *pbx4* function redundantly during early myocardial morphogenesis and outflow tract development. We crossed our *pbx3b*^*scm8*^ strain with the *pbx4*^*b557*^ null allele strain (Kao et al., 2015; Pöpperl et al., 2000) to generate *pbx3b*^*scm8/scm8*^*;pbx4*^*b557/b557*^ embryos. *pbx4*^*b557/b557*^ and *pbx3b*^*scm8/scm8*^*;pbx4*^*b557/b557*^ embryos both show variably disrupted formation of the myocardium and outflow tract (Fig. 1F-G), but *pbx3b*^*scm8/scm8*^*;pbx4*^*b557/b557*^ embryos do not appear to exhibit any more severe myocardial differentiation and outflow tract defects than those that we previously described for *pbx4*^*b557/b557*^ embryos (Fig. 1F-G; Kao et al., 2015). Thus, in zebrafish, *pbx4*, but not *pbx3b*, plays a critical function in early myocardial and outflow tract formation. Studies of gene expression in early human embryos find that *PBX3* is expressed at higher levels than other *PBX* genes (Yan et al., 2013), similar to *pbx4* in zebrafish (Fig. 1C). Functional differences between Pbx genes have been argued to be more likely due to differences in expression than in biochemical activity (Moens and Selleri, 2006; Pöpperl et al., 2000). Therefore, we decided to address the function of the human *PBX3* p.A136V variant in the zebrafish *pbx4* gene.

### Engineering zebrafish *pbx4* p.A131V variant strains

Compared to the human *PBX3* p.A136 site, the zebrafish *pbx4* gene has an orthologous site of A131 (nucleotide C392; Fig. 1A; Fig. 2A). To engineer zebrafish carrying the *pbx4* p.A131V variant, we turned to the CRISPR-Cas9 system (Blackburn et al., 2013; Hwang et al., 2013; Hwang et al., 2015). Although there have been recent improvements, most studies report a low efficiency of introducing precise base changes using CRISPR-Cas9 in zebrafish (Hoshijima et al., 2016; Shah and Moens, 2016; Zhang et al., 2017; Zhang et al., 2018; and references therein), Therefore, to optimize our efforts, we first wanted to ensure that we were using efficient CRISPR-component reagents.

**Figure 2.**
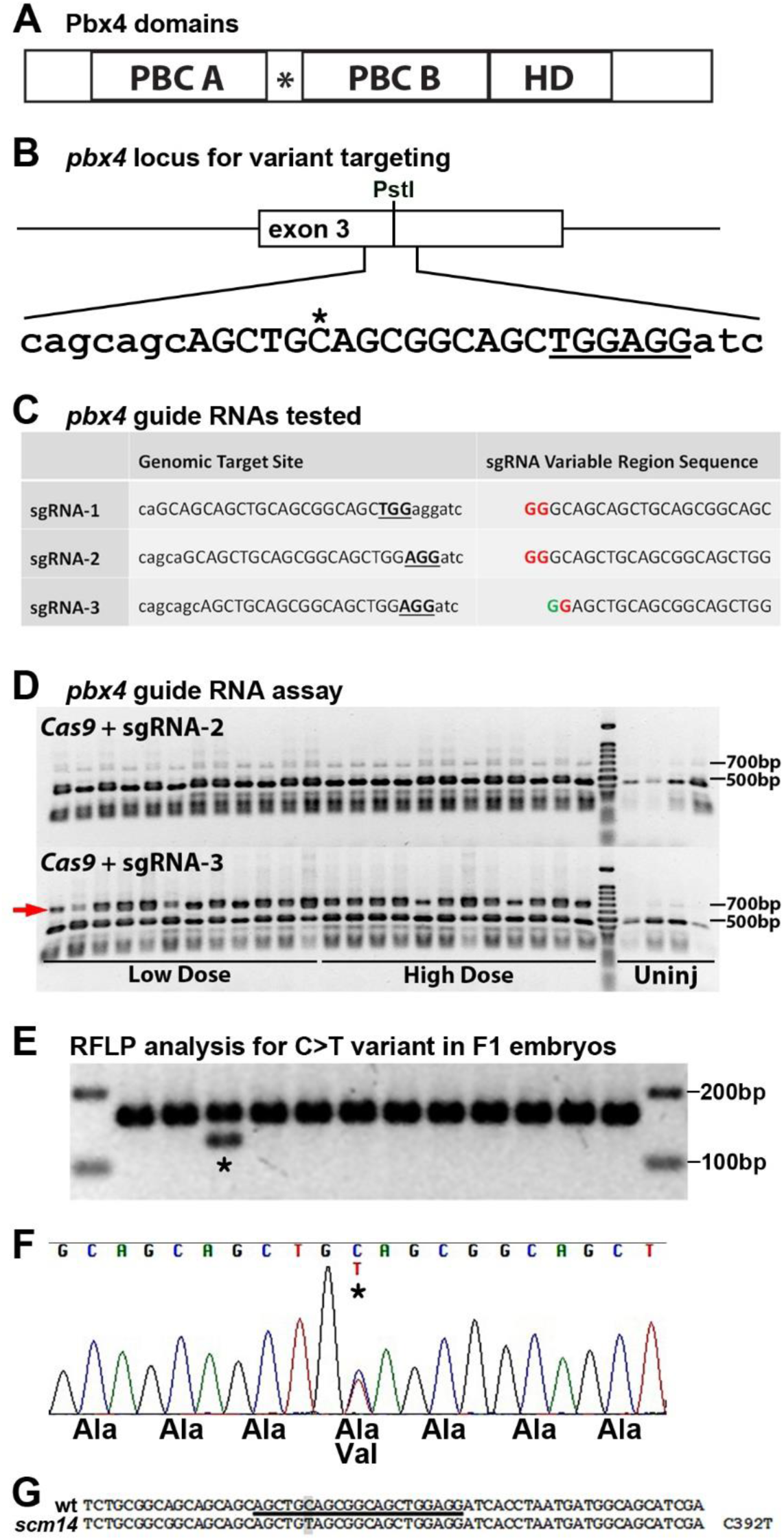
CRISPR engineering of zebrafish *pbx4* p.A131V variant strain. (A) Schematic of zebrafish Pbx4 protein domains. PBC A/B, conserved Pbx domains. *, poly-Alanine tract. HD, homeodomain. (B) *pbx4* locus for targeting the *pbx4 C392/A131* site. The *C nucleotide is nucleotide 392 in the A131 codon. Two adjacent PAM sequences (TGG and AGG) are underlined, and the target site for sgRNA-3 is in uppercase. Surrounding sequence is in lowercase. (C) *pbx4* guide RNAs tested. The middle column shows the genomic region for each guide RNA. The target site is in uppercase. The PAM is in bold underlined uppercase. Surrounding sequence is in lowercase. The right column shows the sequence of the variable region of the sgRNA produced after transcription from pDR274, which begins with a 5’ GG dinucleotide. A red “G” indicates a mis-match with the genomic sequence, while a green “G” indicates a match. (D) DNA restriction fragment length polymorphism (RFLP) analysis of the *pbx4* target region after injection of sgRNA-2 or sgRNA-3. The wild-type PCR amplicon is 684 base pairs (bp) and is cleaved to 241 bp and 443 bp fragments by PstI. Embryos were injected with either a low dose (300pg *Cas9+*12.5pg sgRNA) or a high dose (600pg *Cas9+*25pg sgRNA) of RNAs. In embryos injected with *Cas9+*sgRNA-2, only background levels of non-digested PCR product are present, similar to that seen in uninjected embryos. Injection of *Cas9*+sgRNA-3 results in a large proportion of non-digested PCR product in all embryos assayed (arrow). (E) DNA RFLP analysis of F1 embryos, from CRISPR-Cas9-engineered F0 fish, shows incorporation of the C>T mutation in one embryo (*), due to F0 germline mosaicism. Digestion of the PCR product is dependent on the C>T base change. (F) DNA sequencing chromatogram from heterozygous F1 embryo (* in C) showing precise incorporation of the C>T change. (G) Wildtype (WT) sequence around the *pbx4 C392* site, with the *pbx4* sgRNA target site underlined and C392 highlighted. Sequencing of *scm14* allele shows precise C392T/A131V editing.

We began by designing and testing synthetic guide RNAs. Two adjacent potential PAM sequences (NGG; 5’ PAM and 3’ PAM) are located 3’ of the C392 nucleotide to be changed, so we designed guide RNAs that utilized these PAM sequences and overlapped the C392 nucleotide (Fig. 2B-C). The first guide RNA (sgRNA-1) encompassed the 20 nucleotides (nt) 5’ of the 5’ PAM; the second (sgRNA-2) encompassed the 20 nt 5’ of the 3’ PAM; and the third (sgRNA-3) encompassed the 18 nt 5’ of the 3’ PAM (Fig. 2C). Synthetic single guide RNAs (sgRNA) were produced *in vitro* by annealing pairs of complementary oligos (Table 1) corresponding to the target sequences and ligating them into a plasmid vector containing a T7 promoter and the additional invariant sequences required to form the sgRNA (Hwang et al., 2013). Subsequent transcription of the constructs by T7 polymerase results in the addition of a GG dinucleotide to the 5’ end of the variable region of the sgRNA that binds to the target genomic sequence (Fig. 2C). To test the ability of each sgRNA to induce insertions or deletions (indels) at the target locus, we injected one-cell stage zebrafish embryos with a mixture of *Cas9* mRNA and one of the three sgRNAs, allowed the embryos to develop for 24 hrs, and then collected individual embryos for DNA analysis. Indels were detected by PCR-amplifying the region around the target site and digesting the amplicons with PstI, which has a recognition site within the target site (Fig. 2B,D). We observed a striking difference in the efficacy of the three sgRNAs. The first two sgRNAs produced no or extremely low levels of indels, with 0 out of 24 embryos for sgRNA-1 (not shown) and 0 out of 24 embryos for sgRNA-2 (Fig. 2D) showing a detectable loss of the PstI site. In contrast, 24 out of 24 embryos assayed after injection of *Cas9*+sgRNA-3 had clearly detectable levels of undigested PCR product, indicating disruption of the PstI restriction site by CRISPR-induced indels (Fig. 2D).

**Table 1.**
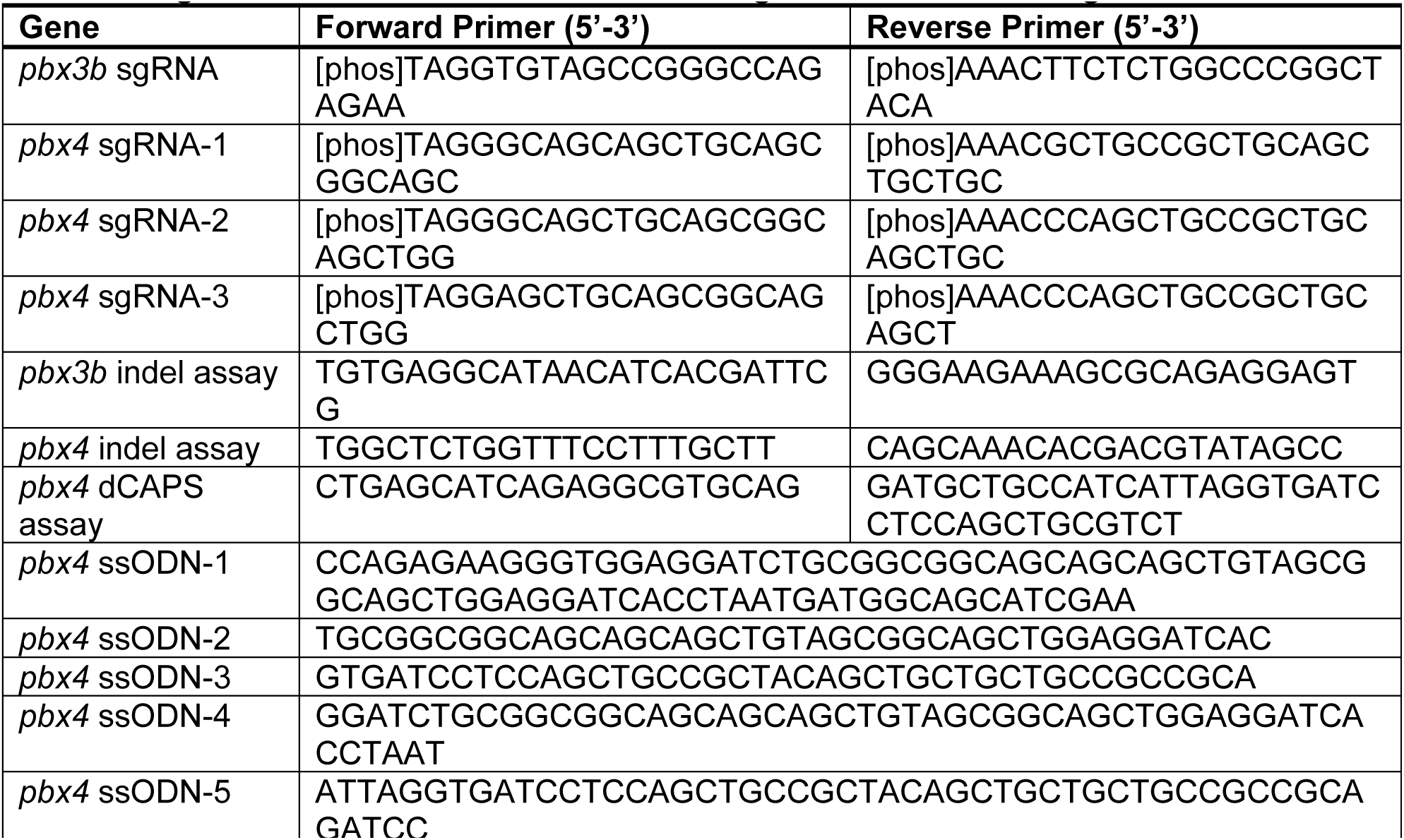
Oligonucleotides for CRISPR-Cas9 mutagenesis and screening

Next, we designed and tested synthetic single-stranded oligodeoxynucleotide donor templates (ssODNs) for introducing the *pbx4* p.A131V variant through homology directed repair (HDR; Hwang et al., 2015). We designed five ssODNs whose sequences overlap the C392 target site but contain a C392T change (corresponding to *C in Fig. 2B; ssODNs listed in Table 1). One ssODN had 40 nt matching the genomic sequence on either side of the mis-matched T and corresponded to the sense strand (ssODN-1). Two oligos had 20 nt homology arms on either side of the base to be changed, with one oligo corresponding to the sense strand (ssODN-2) and the other to the anti-sense strand (ssODN-3). Finally, two oligos had 25 nt homology arms and corresponded to the sense (ssODN-4) and antisense (ssODN-5) strands. We injected zebrafish embryos with *Cas9* mRNA+sgRNA-3+each ssODN, allowed the embryos to develop for 24 hours, and then assayed them for the presence of indels (using the PstI digest, as above for the sgRNA assays) and for the introduction of the C to T change. To detect the specific base change, we designed PCR primers that would create a new AccI restriction site when C392 was changed to a T. Of the five ssODNs tested, two produced detectable levels of the restriction fragment expected from introduction of the C to T base change in some injected embryos (ssODN-4: 5/12 embryos; ssODN-5: 2/12 embryos). The remaining three oligos induced no detectable levels of C to T base change. None of the ssODNs appeared to affect the frequency of induction of indels, with all injection conditions yielding approximately 90% of embryos with some level of indels (similar to that shown in Fig. 2D; data not shown).

Based on these results, we then co-injected the *pbx4* guide RNA (sgRNA-3), *Cas9* mRNA, and the ssODN-4 oligo into 1-cell zebrafish embryos and raised these F0 animals to adulthood. We screened adult F0 fish for those transmitting germline *pbx4* p.A131V mutations by assaying their F1 embryos for incorporation of the C392T mutation, using the same PCR and restriction fragment length polymorphism analysis used in screening the ssODNs (Fig. 2E). Out of 60 F0 adults screened, we identified two independent founder fish transmitting the precise C392T change. We also identified founders carrying a variety of additional alleles (indels) around the *pbx4 (C392)* site (data not shown). We bred F0 fish to wild-type fish to generate F1 fish. We then identified F1 adult fish heterozygous for the precise desired *pbx4* p.A131V variant (Fig. 2F-G). These F1 fish were then bred to wild-type fish to generate a zebrafish strain, *pbx4*^*scm14*^, carrying the *pbx4* p.A131V variant (Fig. 2G). We sequenced the *pbx4* gene in the *pbx4*^*scm14*^ strain to confirm that there were no other changes introduced in *pbx4* (data not shown).

Upon genotyping adult fish derived from a *pbx4*^*scm14/+*^ × *pbx4*^*scm14/+*^ cross, we find the expected frequency of *pbx4*^*scm14/scm14*^ fish (7/32 fish; Chi-square test, P = 0.7788), showing that we obtain viable homozygous adults for the *pbx4* p.A131V variant, while homozygous null *pbx4*^*b557/b557*^ animals die at about 5 days post fertilization (Pöpperl et al., 2000). The adult viability suggests that the *pbx4* p.A131V variant does not, on its own, lead to a significant defect in heart development (as we further confirm below).

To test of the strength of the *pbx4* p.A131V allele, we crossed the *pbx4*^*scm14*^ strain with the null *pbx4*^*b557*^ strain. From a cross of *pbx4*^*scm14/+*^ fish with *pbx4*^*b557/+*^ fish, we find that transheterozygous *pbx4*^*scm14/b557*^ fish survive to adulthood and are present at the expected frequency (10/43 fish; Chi-square test, P = 0.2640). This suggests that the *pbx4* p.A131V allele is likely a weakly hypomorphic allele. We then used the new *pbx4* variant allele strain to test whether the *pbx4 p.A131V* variant functions as a genetic modifier of other CHD genes.

### The *pbx4* p.A131V variant enhances loss of the CHD gene *hand2*

To further test the function of the *pbx4* p.A131V allele, we next wanted to additionally perturb the genetic burden of fish carrying the *pbx4* p.A131V allele. In particular, we decided to directly test whether the *pbx4* p.A131V variant functions as a genetic modifier and enhances the phenotype caused by loss of a known CHD gene, *hand2*. *hand2* encodes a basic helix-loop-helix factor that has critical requirements for embryonic heart development in zebrafish and mice (Srivastava et al., 1997; Yelon et al., 2000), and mutations in the human *HAND2* gene have been associated with CHD (Lu et al., 2016; Shen et al., 2010; Sun et al., 2016; Töpf et al., 2014). In zebrafish, *hand2* is required for both cardiomyocyte differentiation and early myocardial morphogenesis (Garavito et al., 2010; Schoenebeck et al., 2007; Trinh et al., 2005; Yelon et al., 2000;). Furthermore, our previous work used morpholino knockdowns to show that *hand2* and *pbx4* act together in zebrafish early myocardial morphogenesis (Maves et al., 2009). We therefore crossed our *pbx4* p.A131V allele strain (*pbx4*^*scm14*^) with the established null mutant strain, *hand2*^*s6*^ (Yelon et al., 2000). We additionally included the *pbx4*^*b557*^ null allele in these crosses. Fig. 3A shows the genetic crosses used to obtain embryos of all possible homozygous and heterozygous mutant combinations needed for our analyses.

**Figure 3.**
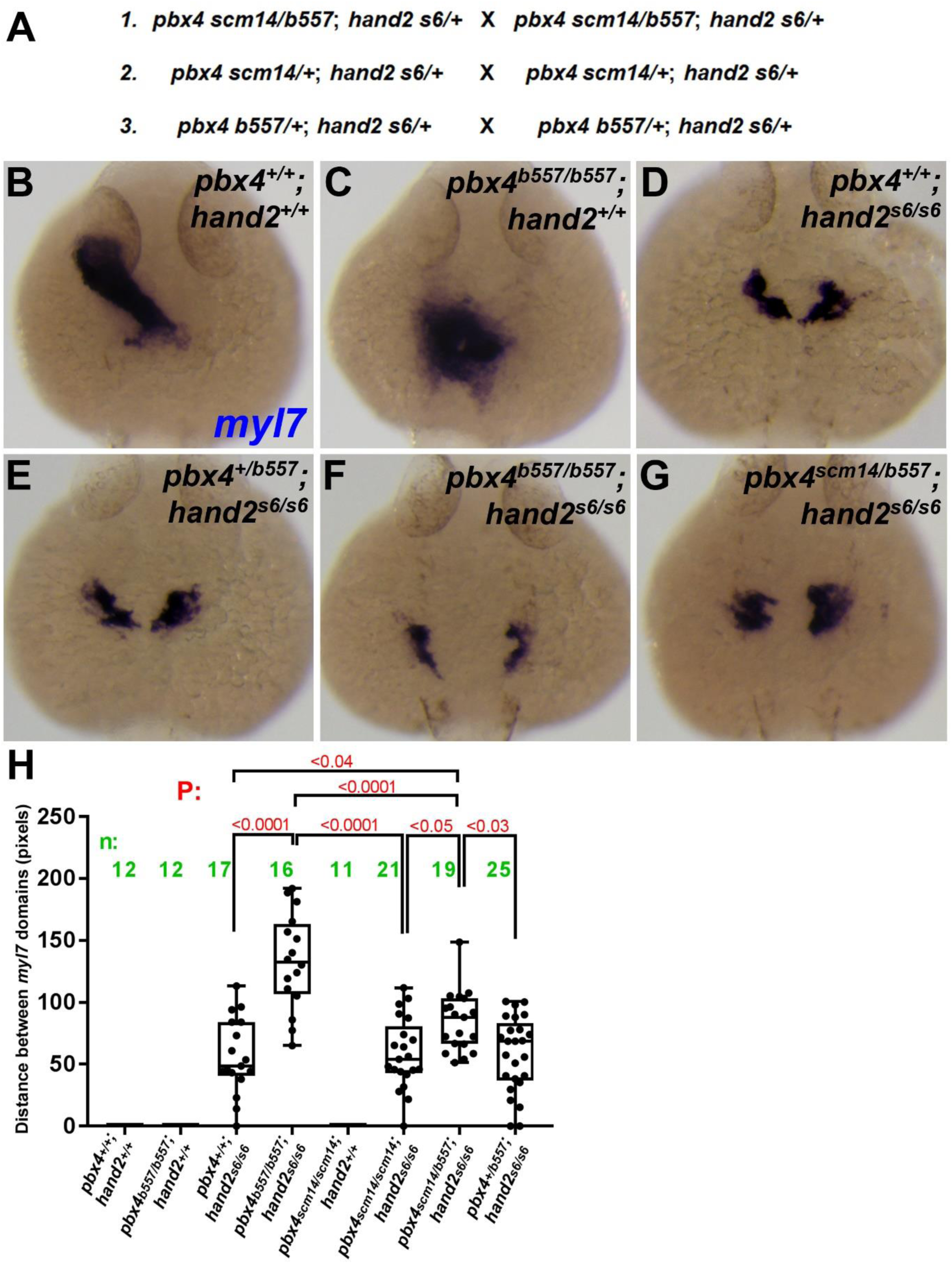
The *pbx4* p.A131V variant enhances the myocardial morphogenesis defect caused by loss of *hand2*. (A) Genetic crosses of zebrafish strains used to obtain embryos for the analyses of *pbx4;hand2* mutant embryos. All adult breeder fish used in crosses 1-3 were “siblings”, derived from the same clutch from a group cross. (B-G) Myocardial marker *myl7* expression at 24 hpf. Dorsal views; anterior is up. (B) The heart tube appears normal in *pbx4*^*+/+*^ *;hand2*^*+/+*^ embryos. (C) *pbx4*^*b557/b557*^*;hand2*^*+/+*^ embryos have a medial heart cone *myl7* domain. (D) *hand2*^*s6/s6*^ embryos show a crescent-shaped myocardial fusion defect of the *myl7* domains, with similar phenotype in *pbx4*^*b557/+*^*;hand2*^*s6/s6*^ (E). (F-G) *myl7* fusion defect is more severe in *pbx4*^*b557/b557*^*;hand2*^*s6/s6*^ and in *pbx4*^*scm14/b557*^*;hand2*^*s6/s6*^. (H) Quantitation of fusion defect of *myl7* domains in different genetic combinations.

We examined early cardiac phenotypes in embryos from these crosses at 24 hpf, using RNA in situ expression of *myl7* to examine myocardial precursors. At 24 hpf, wild-type embryos have formed a heart tube (Fig. 3B; Staudt et al., 2012), and *pbx4*^*scm14/scm14*^ embryos also appear to have a normal heart tube (not shown). As previously described, *pbx4*^*b557/b557*^ embryos show an abnormally-shaped, medially-positioned myocardium (Fig. 3C; Kao et al., 2015), and *hand2*^*s6/s6*^ embryos show reduced, medial, crescent-shaped myocardial domains that have not assembled together properly, with defective fusion of the myocardial precursors at the midline (Fig. 3D; Yelon et al., 2000). We quantified the myocardial fusion defects in these embryos by measuring the distance between the *myl7* domains (Fig. 3H). The *hand2*^*s6/s6*^ myocardial fusion defect is significantly more severe in *pbx4*^*b557/b557*^*;hand2*^*s6/s6*^ embryos, in that the *myl7* domains have greater bilateral separation (Fig. 3F,H). We find that *pbx4*^*scm14/b557*^*;hand2*^*s6/s6*^ embryos also show a more severe effect on myocardial fusion than that seen in *hand2*^*s6/s6*^ embryos, and, critically, the defect is also more severe than that seen in *pbx4*^*b557/+*^*;hand2*^*s6/s6*^ embryos, or in *pbx4*^*scm14/scm14*^*;hand2*^*s6/s6*^ embryos. (Fig. 3E-H). These results show that the *pbx4* p.A131V variant, combined with a null *pbx4* allele, increases the severity, or enhances, the myocardial morphogenesis phenotype caused by loss of *hand2* function. We also analyzed the frequencies of myocardial phenotypes in embryos from these crosses (Table 2). We classified embryos as having a wild-type heart tube, a medial heart cone (the *pbx4*^*b557/b557*^ phenotype), a medial crescent (the *hand2*^*s6/s6*^ phenotype), or a bilateral domain phenotype (the *pbx4*^*b557/b557*^*;hand2*^*s6/s6*^ phenotype). This analysis also shows that the myocardial fusion defect is more severe in *pbx4*^*scm14/b557*^*;hand2*^*s6/s6*^ embryos than in *hand2*^*s6/s6*^ embryos, *pbx4*^*b557/+*^*;hand2*^*s6/s6*^ embryos, or *pbx4*^*scm14/scm14*^*;hand2*^*s6/s6*^ embryos (Table 2). Together, these results demonstrate that the *pbx4* p.A131V variant functions as a genetic modifier and enhances the myocardial morphogenesis phenotype caused by loss of *hand2* function. Therefore, even though the *pbx4 p.A131V* fish are viable and do not have obvious heart defects in an otherwise wild-type genetic background, our results support our hypothesis that the *pbx4 p.A131V* allele can function as a genetic modifier in heart development.

**Table 2.** Genotype-phenotype analysis of *pbx4;hand2* mutant embryos.

| Myocardial Phenotype | *hand2*<sup>+/+</sup> |  |  |  |  |  | *hand2*<sup>s6/+</sup> |  |  |  |  |  | *hand2*<sup>s6/s6</sup> |  |  |  |  |  |
| --- | --- | --- | --- | --- | --- | --- | --- | --- | --- | --- | --- | --- | --- | --- | --- | --- | --- | --- |
|  | *pbx4*<sup>+/+</sup> | *pbx4*<sup>b557/+</sup> | *pbx4*<sup>scm14/+</sup> | *pbx4*<sup>b557/b557</sup> | *pbx4*<sup>scm14/scm14</sup> | *pbx4*<sup>scm14/b557</sup> | *pbx4*<sup>+/+</sup> | *pbx4*<sup>b557/+</sup> | *pbx4*<sup>scm14/+</sup> | *pbx4*<sup>b557/b557</sup> | *pbx4*<sup>scm14/scm14</sup> | *pbx4*<sup>scm14/b557</sup> | *pbx4*<sup>+/+</sup> | *pbx4*<sup>b557/+</sup> | *pbx4*<sup>scm14/+</sup> | *pbx4*<sup>b557/b557</sup> | *pbx4*<sup>scm14/scm14</sup> | *pbx4*<sup>scm14/b557</sup> |
| Wild type heart tube | 14 | 22 | 14 | 0 | 19 | 20 | 35 | 35 | 35 | 0 | 17 | 29 | 0 | 0 | 1 | 0 | 0 | 0 |
| Medial heart cone | 0 | 0 | 0 | 17 | 0 | 1 | 0 | 0 | 0 | 43 | 1 | 5 | 2 | 2 | 3 | 0 | 2 | 0 |
| Medial crescent | 0 | 0 | 0 | 0 | 0 | 0 | 0 | 0 | 0 | 0 | 0 | 0 | 9 | 9 | 12 | 1 | 11 | 2 |
| Bilateral | 0 | 0 | 0 | 0 | 0 | 0 | 0 | 0 | 0 | 0 | 0 | 0 | 6 | 14 | 6 | 15 | 8 | 17 |

Numbers indicate total number of embryos of each genotype with the indicated myocardial phenotype class. Shaded cells indicate genotypes/phenotypes that were compared by Fisher's exact test: *pbx4*<sup>scm14/b557</sup>;*hand2*<sup>s6/s6</sup> versus *pbx4*<sup>+/+</sup>;*hand2*<sup>s6/s6</sup>, P=0.0013; *pbx4*<sup>scm14/b557</sup>;*hand2*<sup>s6/s6</sup> versus *pbx4*<sup>b557/+</sup>;*hand2*<sup>s6/s6</sup>, P=0.021; *pbx4*<sup>scm14/b557</sup>;*hand2*<sup>s6/s6</sup> versus *pbx4*<sup>scm14/scm14</sup>;*hand2*<sup>s6/s6</sup>, P=0.0011.

## Discussion

Here we used CRISPR-Cas9 precision genome editing to successfully engineer a zebrafish strain with a *PBX3* p.A131V variant of unknown significance that was previously identified in a cohort of patients with CHDs. We engineered the human variant into the homologous *pbx4* p.A131V site in zebrafish. We used the engineered zebrafish to demonstrate that the *pbx4* p.A131V allele acts as a genetic enhancer of the known CHD gene *hand2*. Our work underscores the importance of testing the roles of inherited variants as genetic modifiers of CHDs. Our study is the first example, of which we are aware, of using precision genome editing in zebrafish to demonstrate a function for a human disease-associated variant of unknown significance. Our work provides a novel approach toward advancing our understanding of the complex genetics of CHDs and also provides an example for using genome editing in zebrafish to test the causal roles and genetic interactions of human disease-associated DNA variants.

Many approaches in zebrafish and mammalian models have been previously used to characterize functions of human heart disease-associated DNA sequence variants. One study showed that zebrafish embryos could be used to characterize heart function defects caused by a human sodium channel gene *SCN5A* variant, which is strongly associated with human heart disorders such as arrhythmias (Huttner et al., 2013). However, this study employed overexpression of the human gene in transgenic zebrafish. Zebrafish embryos have often been used to test the functions of potential human disease variants through complementation, in which the orthologous zebrafish gene is typically knocked out through genetic mutation or knocked down with antisense morpholinos, and then mRNA injections of a human wild-type or mutant form are used to attempt rescue of a phenotype (Davis et al., 2014). However, due to the inherent transient and mosaic nature of the mRNA injection and overexpression approach, these complementation assays have challenges for examining genetic interactions. A CHD-associated *GATA4* variant has been functionally characterized in a mouse model (Misra et al., 2012), but this was done with conventional embryonic stem cell targeting, which incorporates a targeting vector into the genome. This same *GATA4* variant was also functionally characterized in patient-derived cardiomyocytes induced from pluripotent stem cells, and this analysis provided a deep, systems-level mechanistic understanding of how this particular variant disrupts cardiomyocyte gene expression and function (Ang et al., 2016). However, such cell culture models also have challenges for examining genetic interactions and may not reveal the full effects of a variant on a gene’s function in vivo.

While the development of CRISPR-Cas9 technology has made possible the creation of targeted single-nucleotide changes in zebrafish, the process is still inefficient. Our studies support optimization of the component reagents for successful single-nucleotide editing. We found that different sgRNAs and ssODNs had very different efficacies. Of the three sgRNAs tested, two failed to induce indels at a level we could detect with an RFLP assay, while the third resulted in readily detectable indels in nearly all injected embryos. The highly effective sgRNA has a 20 nt targeting region with a single mis-match at the second position, whereas the ineffective sgRNAs had 22 nt targeting regions and 2 mismatches at their 5’ ends. While sgRNAs with 22 nt protospacers have been used in zebrafish effectively (Hwang et al., 2013), most published studies have used sgRNAs with 20 nt protospacers. One-or two-base mismatches at or near the 5’ end of the protospacer have been shown to be tolerated (Cong et al., 2013; Gagnon et al., 2014; Hwang et al., 2013), and a guanine immediately 5’ to the PAM sequence has been shown to correlate positively with indel frequencies (Farboud and Meyer 2014; Gagnon et al., 2014). Thus, the high incidence of indels seen with *pbx4* sgRNA-3 may result from a combination of optimal protospacer length, the presence of only a single mismatch near the 5’ end of the protospacer, and a G nucleotide in the final position of the protospacer before the PAM. Less is known about the design of homology-directed repair templates to maximize precise editing, and there is conflicting evidence as to whether ssODNs are more efficient at introducing single nucleotide changes than long double-stranded DNA repair templates (Boel et al., 2016; Hoshijima et al., 2016; Hruscha et al., 2013; Hwang et al., 2013; Irion et al., 2014; Liang et al., 2017; Zhang et al., 2018). Previous studies have achieved template-mediated repair using ssODNs of similar length, or longer, to our ssODN-4 (Boel et al., 2016; Hruscha et al., 2013; Hwang et al., 2013; Liang et al., 2017), but homology arms shorter than about 25 nt appear to be ineffective, as we found with our ssODN-2 and ssODN-3.

Our study provides further support for critical roles for PBX-related TALE-class homeodomain transcription factors in heart development and CHDs. Pbx1, Pbx2, and Pbx3 proteins and the Pbx-related factor Meis1 all contribute to outflow tract development in mouse embryos (Chang et al., 2008; Stankunas et al., 2008). Zebrafish *pbx* and *meis* genes are also needed for proper heart development (Guerra et al., 2018; Kao et al., 2015; Maves et al., 2009; Paige et al., 2012). In human cardiomyocyte cell culture models, DNA binding sites for PBX and MEIS factors have been identified as enriched in open chromatin and occurring nearby sites for other cardiac transcription factors (He et al., 2011; Paige et al., 2012; Stergachis et al., 2013; Wamstad et al., 2012). Human sequencing studies have found *PBX* and *MEIS* gene variants associated with CHDs (Arrington et al., 2012; Crowley et al., 2010; Louw et al., 2015; Slavotinek et al., 2017). Notably, a recent study described five patients, each with a CHD and other congenital anomalies, that all had *de novo* missense or nonsense sequence variants in *PBX1* (Slavotinek et al., 2017). The missense *PBX1* variants occurred in or near the homeodomain and likely affect the transcriptional capabilities of PBX1 (Slavotinek et al., 2017). The *PBX3* p.A136V variant that we addressed here is in the more N-terminal poly-Alanine tract, which may bind HDAC proteins (Saleh et al., 2000, although see Choe et al., 2009). The poly-Alanine tract is adjacent to the PBC domains, which interact with Meis and HDAC proteins (Choe et al., 2009; Mann et al., 2009; Saleh et al, 2009). Even though one caveat of our study is that we model the human *PBX3* variant in the zebrafish *pbx4* locus, the poly-Alanine tract is highly conserved in both PBX3 and Pbx4. As future studies identify additional inherited and *de novo* variants in *PBX* genes associated with CHDs, we will gain a better understanding of which human *PBX* genes, and which PBX protein domains, are needed for human heart development.

Our study also provides further support for the oligogenic basis of CHDs. Previous studies of *Pbx* genes in mouse heart development supported a multigenic basis for CHDs (Stankunas et al., 2008). A study that found family members carrying the same *HAND2* mutation, but exhibiting different cardiac defects (Sun et al., 2016), also supports a role for multiple genetic changes influencing the severity of CHDs. In order to reveal a contribution of the *pbx4* p.A131V variant in zebrafish heart development, we had to employ an additional null allele of *pbx4* as well as homozygous loss of *hand2*, conditions that are likely not present in human CHD cases. However, we do not yet have an understanding of the full genetic burden of variants in human CHD cases. A recent study found evidence for rare inherited variants, in genes with a known association with cardiac malformations, in parent-child trios affected by atrioventricular septal defects, and these particular rare variants were generally not observed in control trios (Priest et al., 2016). This study, and our work, support the idea that inherited variants across multiple genetic loci may contribute to the penetrance and expressivity of congenital heart defects, and provide further support to the oligogenic inheritance of CHDs (Fahed et al., 2013; Gelb and Chung, 2014). Forward genetics in mice was used recently to identify an interaction between two genes, *Sap130* and *Pcdha9*, in causing hypoplastic left heart syndrome (HLHS; Liu et al., 2017). Intriguingly, this study also identified one human HLHS subject with variants in both *SAP130* and *PCDHA13*, a *Pcdha9* homolog (Liu et al., 2017). In mice, *Sap130* and *Pcdha9* may work combinatorially, each regulating different pathways important for proper heart development (Liu et al., 2017). Although we previously described *pbx4* and *hand2* as working together in zebrafish heart development (Maves et al., 2009), it is possible that they are also working through parallel pathways. Future studies that continue to take advantage of animal models, as well as human genetic sequence data, will further enhance our understanding of the complex genetics of human diseases such as CHD.

## Materials and Methods

### Zebrafish husbandry

All experiments involving live zebrafish (*Danio rerio*) were carried out in compliance with Seattle Children’s Research Institute IACUC guidelines. Zebrafish were raised and staged as previously described (Westerfield, 2000). Staging time refers to hours post fertilization (hpf) at 28.5 °C. The wild-type stock and genetic background used was AB. The *pbx4*^*b557*^ mutant strain was previously described and is likely a null allele (Pöpperl et al., 2000). *pbx4*^*b557*^ genotyping was performed as previously described (Kao et al., 2015) using forward primer 5’ACTCGGCGGACTCTCGCAAGC3’ and reverse primer 5’GGCTCTCGTCGGTGATGGCCATGATCT3’. The genotyping PCR product is 128 base pairs; digesting with XbaI yields a 98 base pair product from the mutant allele. In some cases, *pbx4*^*b557*^ animals were genotyped using a KASP assay (LGC Genomics). Reactions were run and fluorescence was measured in a Bio-Rad CFX96, and genotypes were assigned using the Bio-Rad CFX Manager software, according to instructions provided by LGC. Details for ordering the *pbx4*^*b557*^ KASP assay are available upon request. The *hand2*^*s6*^ mutant strain was previously described and is a deletion of the *hand2* locus (Yelon et al., 2000). For *hand2*^*s6*^ genotyping, we devised a qPCR assay for the number of copies of the *hand2* gene. *hand2* was amplified with forward primer 5’ACCAAAGCGTACTCCGTCTG3’ and reverse primer 5’CAGCGAAGGAATAGCCGTCA3’. *pbx4* was also amplified as a normalization control using forward primer 5’GCCGTTAAAACAGCCGTGG3’ and reverse primer 5’GTGTTGCTGGAGAGTTTGCC3’. Reactions were run in duplicate for both genes using the KAPA SYBR FAST kit (KAPA Biosystems KK4600) on a Bio-Rad CFX96 machine, the resulting Ct values were averaged, and a ΔCt was calculated to distinguish *hand2* homozygous, heterozygous, and wild-type embryos.

### Pbx phylogenetic analysis

Accession numbers for the sequences aligned in Figure 1A are given in Table 3. Sequences were aligned using Clustal Omega (www.EMBL.org). The phylogeny was constructed with PhyML using the following settings: Substitution Model LG; Tree Improvement SPR & NNI; Bootstrapping 500 (www.ATGC-montpellier.fr). The phylogenetic tree (Figure 1B) was drawn with TreeDyn (www.phylogeny.lirmm.fr).

**Table 3.** Pbx accession numbers

| **Pbx gene** | **Protein sequence** |
| --- | --- |
| Human *PBX1* | NP_002576.1 |
| Mouse *Pbx1* | NP_001278437.1 |
| Zebrafish *pbx1a* | NP_571689.1 |
| Zebrafish *pbx1b* | NP_001077322.2 |
| Human *PBX2* | NP_002577.2 |
| Mouse *Pbx2* | NP_059491.1 |
| Zebrafish *pbx2* | NP_571687.1 |
| Human *PBX3* | NP_006186.1 |
| Mouse *Pbx3* | NP_058048.1 |
| Zebrafish *pbx3a* | XP_005171794.1 |
| Zebrafish *pbx3b* | CAQ13247.1 |
| Human *PBX4* | NP_079521.1 |
| Mouse *Pbx4* | NP_001020125.1 |
| Zebrafish *pbx4* | NP_571522.1 |
| Fruit fly *exd* | NP_523360.1 |

### qRT-PCR

Total RNA was isolated using TriZol (Ambion; Thermo Fisher Scientific) and reverse-transcribed with the SensiFAST cDNA Synthesis kit (Bioline BIO-65053). Primers were designed using Primer-BLAST such that they either span an intron or one of the pair spans an exon-exon boundary. Primers are listed in Table 4. Real-time PCR was carried out using the KAPA SYBR FAST kit (KAPA Biosystems KK4600) on a Bio-Rad CFX96 machine. Ct values for the zebrafish *pbx* genes were corrected for observed primer efficiencies and normalized to *odc1*.

**Table 4.**
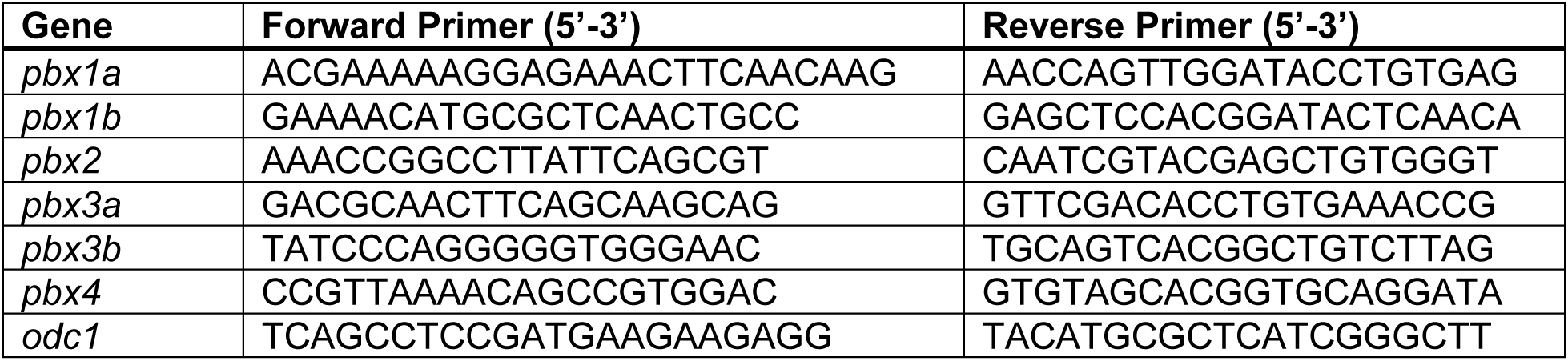
Oligonucleotides for qRT – PCR

### Generation of mutant zebrafish strains with CRISPR-Cas9

Single guide constructs were made by annealing pairs of oligos (listed in Table 1) and ligating them into BsaI-digested pDR274 (Hwang et al., 2013). The single guide plasmids were digested with DraI, and guide RNA was transcribed with the T7 Maxiscript kit (Ambion; Thermo Fisher Scientific). *Cas9* mRNA was made by transcribing PmeI-digested pMLM3613 (Hwang et al., 2013) with the T7 Ultra kit (Ambion; Thermo Fisher Scientific). One-cell-stage zebrafish embryos were injected with 600pg (for *pbx3b*) or 400pg (for *pbx4*) of *Cas9* mRNA and 25pg of a single guide RNA in a volume of 2nL. To generate the desired single nucleotide change in *pbx4*, 50pg of an ssODN (Table 1) was co-injected with *Cas9* mRNA and the single guide RNA. Zebrafish were screened for Cas9-generated mutations (insertions or deletions, “indels”) by amplifying with primers flanking the target site (indel assay primers, Table 1) followed by a digest to test for loss of a restriction site just upstream of or overlapping the PAM site (HpyAV for *pbx3b*, PstI for *pbx4*). To detect the single-nucleotide change in *pbx4*, a dCAPS assay was designed (http://helix.wustl.edu/dcaps/; Neff et al., 1998) such that PCR amplification of the mutant allele generated an AccI site (*pbx4* dCAPS assay primers in Table 1). Mutant alleles were identified in heterozygous F1 or F2 fish by Sanger sequencing and deconvolving the chromatograms with Poly Peak Parser (http://yosttools.genetics.utah.edu/PolyPeakParser/).

### Whole-mount RNA in situ hybridization and measurement analysis

The following cDNA probes were used: *myl7* (Yelon et al., 1999) and *elnb* (Miao et al., 2007). Whole-mount *in situ* hybridization colorimetric and fluorescent *in situ* staining was performed as previously described (Maves et al., 2007; Talbot et al., 2010), with the following modifications. Embryos at 60 hpf were depigmented in 1 part 0.1% KOH (vol.): 1 part 1X PBS-0.1% Tween (vol.): 0.1 part 30% hydrogen peroxide (vol.) for 2 hours at room temperature with gentle agitation. Hybridizations for both colorimetric and fluorescent *in situ* experiments were performed in hybridization buffer with 5% dextran sulfate. Following staining, tail clips from post-*in situ* hybridized embryos were lysed and genotyped for *pbx3b*^*scm8*^, *pbx4*^*b557*^, *pbx4*^*scm14*^, and *hand2*^*s6*^ as above. Fluorescent *in situs* were imaged on a Leica SP5 confocal with a 40x water immersion objective.

For *myl7* measurements, 24 hpf embryos stained for *myl7* were individually imaged at a consistent magnification on a stereomicroscope. ImageJ was used to measure the area of *myl7* expression and the distance between bilateral domains of expression, expressed in pixels. For embryos with separate bilateral *myl7* domains, three distance measurements were made for each embryo: between the medial edges of expression at the anterior extent of expression, between the medial edges of expression at the posterior extent of expression, and at the approximate midpoint between the anterior and posterior measurements. These three values were then averaged. For embryos with a single midline domain of *myl7* expression, a distance value of 0 was assigned. For embryos with a crescent-shaped domain, the posterior distance measurement was 0 and the middle and anterior measurements were made as for separate bilateral domains. The averages for the *myl7* distances among the different genotypes were compared using one-way ANOVA and p-values were corrected for multiple comparisons using Tukey’s test. ANOVA was performed and the box plots were made in GraphPad Prism 7. The boxes extend from the 25th to 75th percentiles, the whiskers are at the minimum and maximum, and the bar within the box represents the median.

## Acknowledgements

We thank Fred Keeley and Debbie Yelon for generously providing reagents. We gratefully acknowledge advice from Stephen Ekker and Cecilia Moens on CRISPR editing in zebrafish. We thank Jerry Ament, Dante D’India, and the SCRI Aquatics Facility staff for expert zebrafish maintenance.

## Author contributions statement

G.H.F.III, K.I., D.P., and L.M. conducted experiments, G.H.F.III and L.M. designed experiments, and G.H.F.III and L.M. wrote the paper with input from all authors.

